# Single-molecule variation in telomeric sequence and structure across humans

**DOI:** 10.64898/2026.05.01.722200

**Authors:** Danilo Dubocanin, Mitchell R. Vollger, Shane J. Neph, Maria Sara Del Rio Pisula, Julian K Lucas, Adriana E Sedeño-Cortés, Ben J Mallory, Taylor D Real, Human Pangenome Reference Consortium, Floris P. Barthel, Nicolas Altemose, Andrew B. Stergachis

## Abstract

The repetitive architectures of telomeric and subtelomeric regions have obscured studies of their genetic variation and chromatin organization across the human population. Here, we integrate near-complete diploid genome assemblies from 212 individuals with matched long-read sequencing data to construct an atlas of 316,146 telomere-spanning molecules across 12,080 chromosome-end-resolved telomere arrays. This atlas reveals that nearly every chromosome end harbors a structured and unique pattern of telomere variant repeats (TVR), or TVR code, with subtelomere-proximal TVR codes being heritable, somatically stable, and influenced by subtelomeric TAR1 regulatory elements. Despite ongoing cycles of telomere shortening and elongation in the germline, proximal TVR codes are maintained across the human population. These TVR codes expose rare telomerase-independent events that lengthen telomeres in the germline, including interchromosomal telomere exchange and recurrent internal duplications within telomere arrays. Furthermore, single-molecule chromatin fiber sequencing across 26,972 molecules spanning the telomere-subtelomere boundary confirms that TVR-rich regions adopt telomeric chromatin but introduce discrete discontinuities into otherwise compact telomeric chromatin fibers. Together, our results link chromosome-end sequence variation to telomere cap formation and telomerase-independent telomere extension mechanisms in the human germline.

## Introduction

In humans, telomeres consist of tandem TTAGGG repeats^1^ bound by the shelterin complex and associated chromatin proteins that together form a protective telomere cap^2^. Disruption of telomere integrity leads to genome instability and contributes to aging and cancer^3^, highlighting the importance of understanding how telomere structure is established and maintained.

Human telomeres are often depicted as homogeneous arrays of TTAGGG repeats, and most studies to date have focused on telomeric length rather than sequence composition^4–7^. However, the proximal regions of many telomeres contain telomere variant repeats (TVRs), degenerate repeats that deviate from the canonical hexameric motif^8–14^. Most previous analyses of TVRs have been limited to small numbers of chromosome ends or individual telomeres, leaving the broader population-scale structure of TVRs largely unexplored. Furthermore, given the divergence of TVRs from the canonical TTAGGG telomere repeat, it has remained unclear whether TVR regions along telomere arrays form chromatin consistent with the telomere cap^14^. As a result, fundamental questions remain unresolved: whether TVR patterns represent stable features of chromosome ends or are continually erased in the germline during cycles of telomere shortening and extension, how telomeric sequence diversity evolves across individuals and generations, and what impact TVRs have on the structure and function of the overlying telomere cap.

Telomeres also exist within a specialized subtelomeric regulatory environment that includes Telomere Associated Repeat 1 (TAR1) elements capable of influencing telomere chromatin and transcription^15^. Specifically, TAR1 elements drive transcription of telomeric repeat-containing RNA (TERRA)^16,17^, a long non-coding RNA implicated in telomere regulation and chromatin organization. In addition, TAR1 elements harbor CTCF-binding sites that may shape local chromatin structure^15,18,19^. However, because telomeres and subtelomeres are highly repetitive, their chromatin organization has been difficult to resolve at high resolution. Consequently, it remains unclear how subtelomeric regulatory elements are organized at the level of individual molecules, how their chromatin states vary across cell types, and how this regulatory architecture interfaces with the underlying telomeric repeat array to influence telomere stability and function.

Here we combine near-complete genome assemblies^20^ with long-read sequencing and single-molecule chromatin profiling^21^ to generate a population-scale atlas of human telomere sequence and chromatin structure. We show that nearly every chromosome end contains a structured pattern of TVRs that form a telomere-specific TVR code. These TVR codes are incorporated into the telomere cap chromatin structure, and reveal principles of telomere inheritance, recombination, and telomerase-independent extension, and demonstrate that telomeric sequence variation and subtelomeric regulatory elements jointly shape the chromatin organization and sequence dynamics of human chromosome ends.

## Results

### An atlas of single-molecule human telomeres

Telomeres are among the most repetitive and length-polymorphic regions of the human genome, and their adjacent subtelomeres contain repeat elements and undergo frequent genomic rearrangements, all of which can challenge existing read mapping strategies for chromosome end-specific telomere studies. To quantify the extent of this challenge, we initially evaluated a sample containing long read PacBio HiFi data and a near telomere-to-telomere (T2T) diploid assembly generated as part of HPRC (HG00099). We observed that HG00099 PacBio HiFi reads mapped to the CHM13 haploid reference genome^22^ resulted in uneven coverage across the 48 telomere-subtelomere junctions, with reads spanning these regions frequently exhibiting numerous base mismatches and low mapping quality (**Extended Data Fig. S1a**). Mapping the same reads to the HG002 T2T diploid genome benchmark assembly^23^ did not resolve these issues, underscoring the limitations of using a fixed reference genome to study highly repetitive telomeric DNA (**Extended Data Fig. S1a,b**). In contrast, these artifacts were largely eliminated when reads were mapped to the donor-matched HG00099 diploid near-T2T assembly^24^, demonstrating that donor-specific assemblies enable the more accurate mapping of reads to telomere-subtelomere junctions (**Extended Data Fig. S1a,b**). However, we observed that even though donor-matched assemblies more accurately capture subtelomeric sequence, substantial differences remained within the telomere arrays between the donor-matched assemblies and the single-molecule reads from that donor in both sequence composition and structural organization (**Extended Data Fig. S1c**), highlighting that telomere arrays represent an unsolved challenge with current genome assembly algorithms, and the importance of evaluating telomere sequences directly from individual sequencing reads from a donor.

Based on these observations, we constructed a chromosome-end-resolved atlas of human telomeres by integrating 212 diploid near-T2T assemblies from the Human Pangenome Reference Consortium (HPRC) release 2 with matched long-read datasets from the same individuals, including 212 PacBio HiFi and 51 Oxford Nanopore Technologies (ONT) R10 datasets. Mapping reads to each donor’s own assembly identified 316,146 telomere-spanning molecules (218,015 PacBio; 98,131 ONT) across 12,080 chromosome-end telomere arrays, with a median of 12 (PacBio) and 24 (ONT) reads per telomere array (**Extended Data Fig. S2**). Using donor-specific assemblies minimized reference bias in telomere sequence composition and enabled accurate assignment of chromosome origin, including for telomeres adjacent to segmental duplications and acrocentric chromosome arms. This chromosome-end-resolved dataset provides a population-scale framework for studying telomere sequence diversity and establishes the foundation for examining how telomeric sequence variation arises and is maintained.

### Human telomeres harbor structured TVR codes

Having established a chromosome-end-resolved telomere atlas, we next sought to determine how telomeric sequence varies along individual telomere arrays. To characterize telomeric sequence composition across all 316,146 telomere-spanning molecules, we first anchored each read at its telomere-subtelomere boundary, enabling reference-free evaluation of the telomeric DNA portion of each read at single-molecule resolution (**Fig. 1a**). We next identified all telomeric bases that could not be assigned to canonical TTAGGG repeats, generating a per-molecule atlas of TVR sequence architecture (**Fig. 1b**). Comparison of multiple molecules originating from the same chromosome end revealed highly consistent TVR patterns within individuals (**Fig. 1b-d, Extended Data Fig. S3a,b**). Moreover, TVR patterns derived independently from PacBio and ONT sequencing were highly concordant (**Fig. 1c**; Pearson’s r = 0.929, **Extended Data Fig. S3a**), confirming both the reproducibility of TVR architecture and the accuracy of TVR detection across sequencing platforms. Intra-individual single-molecule variability was most apparent among TVRs located distally from the subtelomere-telomere boundary^14,25^, consistent with somatic telomere shortening into the TVR array followed by somatic telomerase-mediated re-extension^9,26,27^ (**Extended Data Fig. S4**). Nevertheless, the overall reproducibility of proximal TVR architecture across molecules indicates that most of the telomere array is not continuously erased and rebuilt, but instead is maintained, resulting in stable chromosome-end-specific sequence patterns within individuals.

**Fig. 1.**
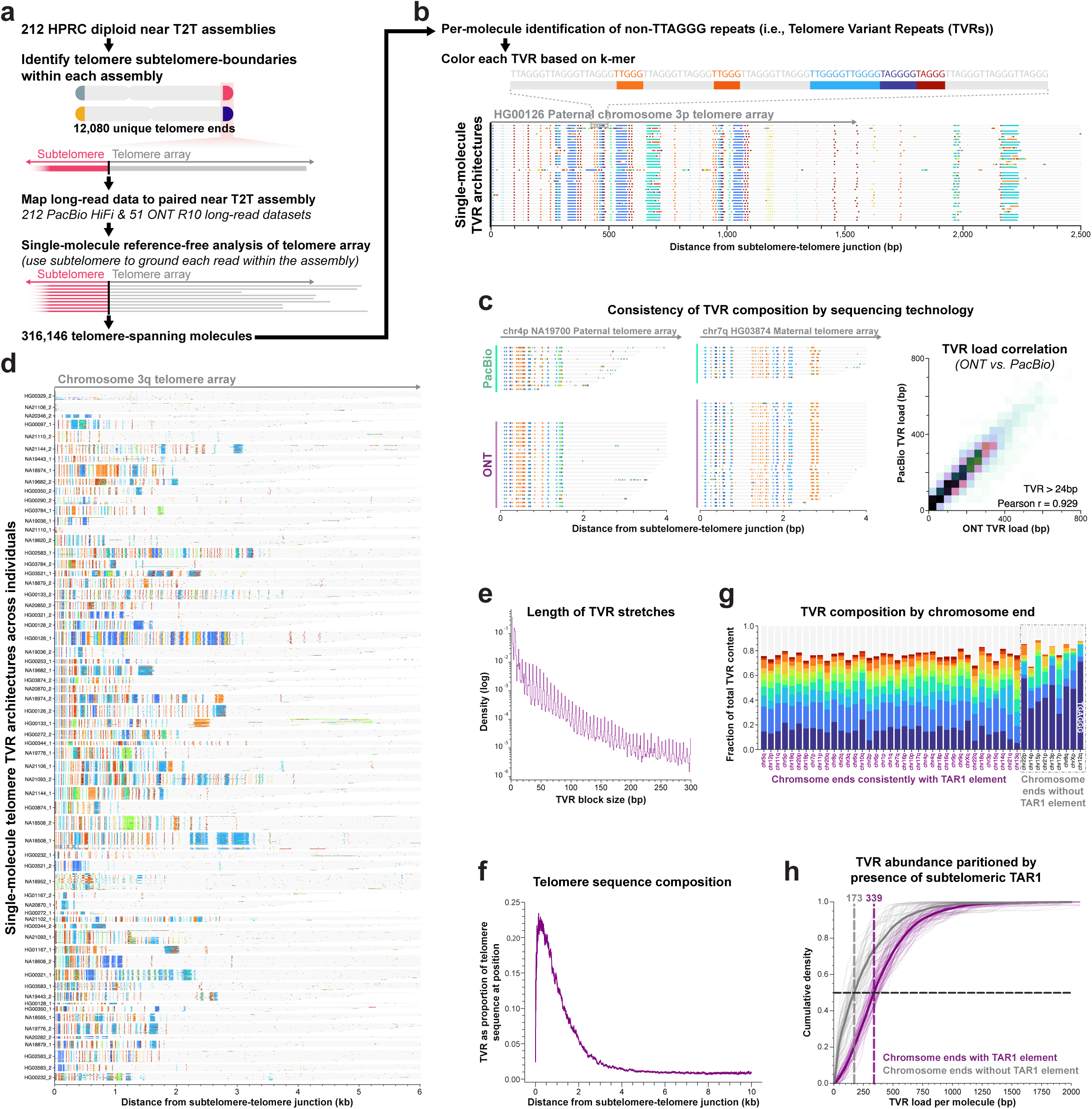
Telomere Variant Repeats detected across HPRC samples. **a**, Overview of the workflow for identifying TVRs across HPRC long-read datasets. **b**, Example of TVR calls along single molecules in HG00126 at maternal chr3p using ONT R10 data. Each row represents a single read anchored at the subtelomere boundary (0bp): gray indicates canonical TTAGGG repeats, and color denotes any deviation from the TTAGGG array (Methods, TVRs over 300bp are not shown). Reads within each haplotype sorted by the length of their telomeric array segment. **c**, Comparison of single-molecule TVR calls between ONT R10 and PacBio HiFi data. Single-molecule plots (left and middle) show TVRs ranging from 6 bp to 300 bp in size. The 2D histogram (right) displays the TVR load correlation for chromosome ends with a minimum of 10 ONT R10 reads and 5 PacBio HiFi reads. Load is calculated as the mean TVR amount (in bp) per chromosome end per haplotype for each technology across molecules greater than 1 kb in length (300bp ≥ TVR sizes ≥ 24bp). **d**, TVRs across single molecules from all chr3q ends in the HPRC dataset (ONT R10). Each row represents a single read, with gray indicating TTAGGG repeats and color representing specific TVRs according to a shared color map. Data is filtered to include a minimum of 5 reads per individual haplotype end, with TVR sizes ranging from 1 to 300 bp. **e**, Distribution of TVR block sizes across all ONT R10 molecules with at least 1 kb of telomeric sequence. **f**, TVRs as a proportion of the telomeric sequence per position for ONT R10 reads with at least 1 kb of telomeric sequence. **g**, Relative distribution of the 10 most common TVR 6-mers total (>6 bp), shown at each chromosome end. TVRs larger than 6 bp are partitioned into consecutive 6-bp windows, advancing with a 6-bp step size until the window passes the end of the TVR. **h**, Cumulative distribution of the TVR amount across molecules, partitioned by whether the corresponding chromosome arm contains a TAR1 element within 10 kb of the subtelomere–telomere boundary. Each faint line represents a single individual, bold line is the mean distribution across samples.

Across chromosome ends, telomere arrays varied markedly in both the abundance and sequence composition of TVRs (**Fig. 1d, Extended Data Fig. S3b**). Most TVRs occurred as 6-mers, with the G-rich repeat TTGGGG, which is also the tetrahymena telomere sequence^28^, representing the most frequent variant (**Fig 1e, Extended Data Fig. S3c-e**). Notably, when multiple TVRs occurred in tandem, they typically consisted of identical 6-mer sequences (**Fig 1d, Extended Data Fig. S3b,d**), indicating that TVR formation is non-random. TVRs were strongly biased toward the subtelomere-proximal region of the telomere array, with 49.7% located within the first 1 kb and 94.5% within the first 5 kb (TVRs > 24bp, ONT R10) despite a mean telomeric read length of 6.98 kb (**Fig. 1f; Extended Data Fig. S3f, S5 ;** ONT read length, mean PacBio read length of 5.7 kb). In contrast, telomere arrays from nine chromosome ends were consistently enriched across donors for one specific TVR sequence, TGAGGG^9^, and harbored substantially fewer overall TVRs (**Fig. 1g; Extended Data Fig. S6**). Together, these observations demonstrate that nearly every human chromosome end harbors a distinct, structured “TVR code”^11,14^, suggesting that telomeric sequence variation is shaped by chromosome-end-specific constraints rather than random mutational processes.

### TVR codes are influenced by subtelomeric TAR1 regulatory elements

The strong chromosome-end specificity of TVR patterns suggested that local subtelomeric architecture might influence telomere sequence composition. Annotation of subtelomeric sequence across all telomere arrays revealed that most chromosome ends are directly adjacent to TAR1 elements^29^, which encode the TERRA promoter and clustered CTCF-binding sites (**Extended Data Fig. 7a,b**). TAR1 elements are frequently absent from acrocentric short arms (13p, 14p, 15p, 21p, and 22p) as well as 3p and 17p, and are consistently absent from chromosome arms 8q, 12q, and Xp, while being nearly ubiquitous at other chromosome ends (**Extended Data Fig. 7a,b**). Telomere arrays adjacent to TAR1 elements exhibited a substantially increased TVR burden, with a 96% higher median TVR load than TAR1-lacking telomeres (**Fig. 1h; Extended Data Fig. S7c**). Moreover, when TVRs were present on TAR1-deficient telomeres, they were disproportionately composed of the TGAGGG 6-mer variant (**Fig 1g, Extended Data Fig. S6**). The enrichment of TVRs at TAR1-containing chromosome ends, coupled with their depletion and altered sequence composition at TAR1-deficient ends, implicate TAR1 elements as key cis determinants of telomere sequence architecture.

We next investigated the architecture and cell-type dynamics of TAR1 element-encoded regulatory features using Fiber-seq data from two developmentally distinct cell lines with paired T2T assemblies (*i.e.,* HG002 lymphoblastoid cells and CHM13 hTERT-immortalized hydatidiform mole cells)^22,23,30^. Fiber-seq uses a non-specific adenine methyltransferase (MTase) to simultaneously capture native DNA sequence and overlying DNA-binding protein footprints with single-nucleotide and single-molecule precision^21^. Furthermore, as Fiber-seq is grounded in long-read sequencing, it can be readily mapped to complex genomic loci, such as telomeres^25,26,30^. We observed that within HG002 cells TAR1 elements were predominantly occupied by regularly spaced nucleosome arrays and extended MTase-protected regions, consistent with a largely closed chromatin state. In contrast, in CHM13 cells, TAR1 elements tended to adopt a prominent accessible chromatin domain immediately proximal to the telomere-subtelomere boundary (**Extended Data Fig. S8a,b**). These accessible regions encompassed the subtelomeric TERRA promoter as well as the clustered TAR1 CTCF-binding sites, and displayed both inter-telomere and intra-telomere variability in accessibility (**Extended Data Fig. S8a,b**). Furthermore, within TAR1 elements that had chromatin accessibility, CTCF-binding sites showed robust single-molecule occupancy in a highly stereotyped orientation, with nearly all motifs positioned such that the CTCF N-terminus faced away from the telomere, creating a structural boundary between the subtelomere and the TERRA promoter and telomere array (**Extended Data Fig. S8c-g**). Notably, CHM13 chromosome ends lacking TAR1 uniformly lacked this accessible domain, indicating that TAR1 elements are necessary, though not alone sufficient, to establish this subtelomeric regulatory architecture (**Extended Data Fig S8b**). Together, these findings reveal that TAR1 elements form a cell-selective regulatory module at human chromosome ends that is associated with telomere TVR abundance and sequence composition.

### Ancestral TVR codes are stable across generations through the human germline

Although telomeres are often described as highly dynamic due to cycles of shortening and telomerase-mediated re-extension, select chromosome end TVR codes have been shown to be stable through single parent-offspring transmission events^9,14^(**Extended Data Fig. S9**). Telomere lengths show a heritability estimate of 0.7, indicating that telomere lengths can be maintained across generations^9,14,31–33^, or are quickly reshaped during telomere maintenance in the germline. Given our finding that each chromosome-end telomere array harbors a largely distinct TVR code, we asked whether identical or highly similar TVR codes could be detected across chromosome ends from different individuals, consistent with descent from a common ancestral telomere. To quantify similarity among TVR codes, we established a consensus TVR sequence for each chromosome-end telomere array using a customized alignment algorithm for TVRs^34^ optimized for pairwise comparison of telomere arrays containing at least five telomeric molecules (**Fig. 2a**, Methods). Using these alignments, we measured similarity between TVR codes from all pairs of telomere arrays as a function of distance from the telomere-subtelomere boundary (**Fig. 2b**). At a similarity threshold of 80%, we found that 32% (3,032 of 9,489) of telomere arrays contain a proximal TVR code ≥200 bp that is present in multiple individuals (**Fig. 2c,d,e; Extended Data Fig. S10, S11**, PacBio). Importantly, these shared TVR codes were strongly localized to the subtelomere-proximal region of the telomere array, with similarity declining sharply with increasing distance into the telomeric repeat (**Fig. 2c,e; Extended Data Fig. S10, S11**). Specifically, only 0.13% of telomere arrays contained a proximal TVR code ≥1,000 bp in length that is present in multiple individuals (12 of 9,489; **Fig. 2e**). This rapid positional decay suggests that proximal TVRs are preferentially immune from being progressively remodeled during telomere maintenance in the germline.

**Fig. 2.**
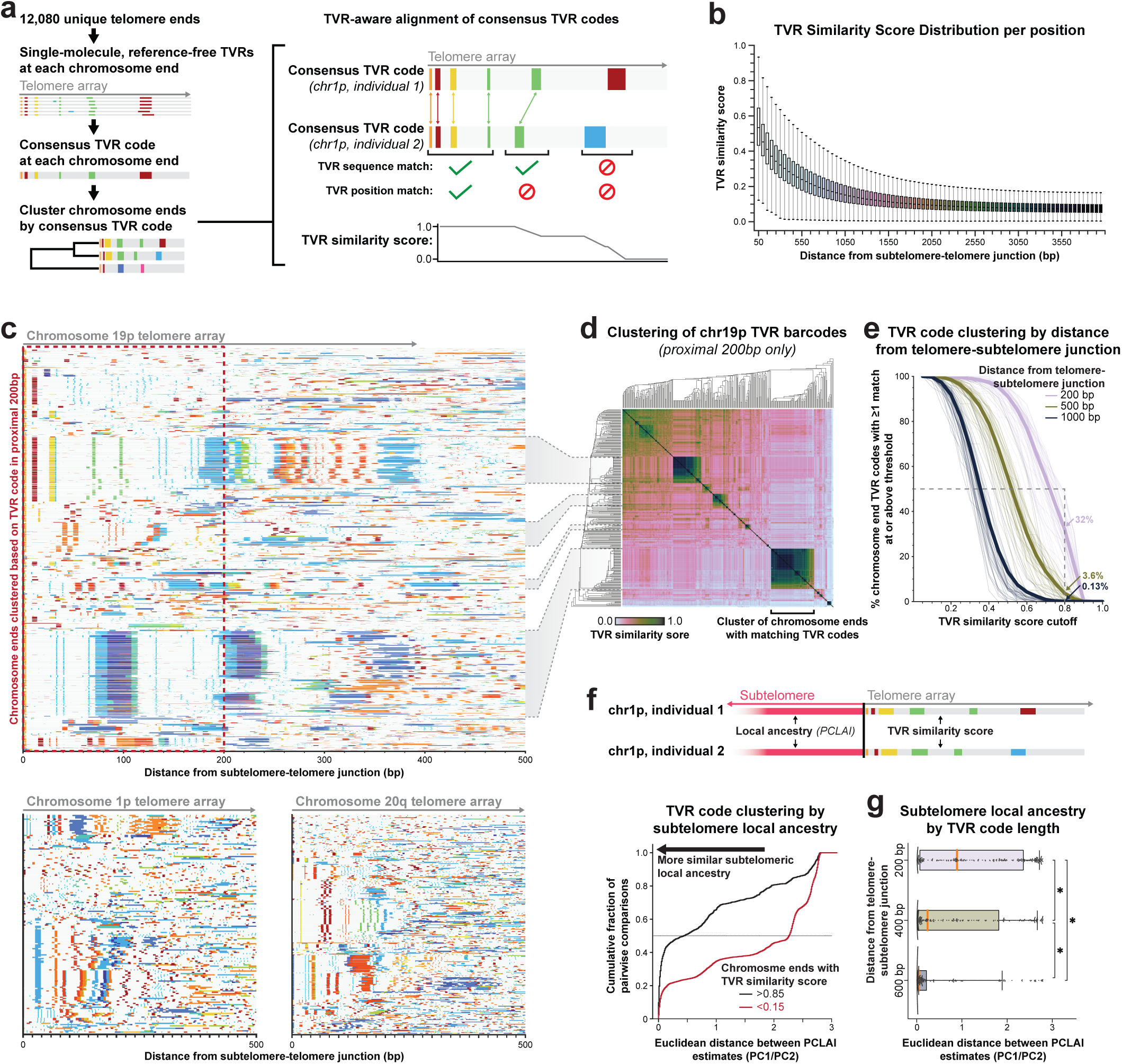
Ancestral TVR codes are stable across generations through the human germline. **a**, Overview of TVR consensus generation and mapping to generate TVR similarity scores (Methods). **b**, Distribution of TVR similarity scores at various distances from the subtelomere. Note that consensus generation lacks an alignment step, thus quality degrades with telomeric depth as stochastic read errors shift relative TVR positions. **c**, Top, PacBio HiFi single-molecule TVR plots of the subtelomere-proximal 500bp of telomeric sequence in chr19p. Each row is a single-molecule, grouped by individual haplotype, and clustered based on the proximal 200bp consensus TVR code (*n =* 304 haplotypes). Bottom, Randomly sampled HiFi molecules per haplotype, clustered based on the proximal 200bp consensus TVR code (chr1p *n =* 175 haplotypes; chr20q *n =* 303 haplotypes). **d**, Dendrogram and corresponding pairwise heatmap clustered based on TVR consensus similarity scores in the proximal 200bp of chr19p corresponding to **c** (top). **e**, Proportion of chromosome-end TVR codes shared with at least one other chromosome end across various similarity thresholds (0.05 increments). Faint lines represent individual chromosome arms; bold lines indicate proportion across all arms. **f**, (top) Schematic describing how local ancestry and TVR similarity scores are partitioned across subtelomere/telomere boundaries. (bottom) Cumulative distribution of the mean Euclidean Distance of PCLAI in the terminal 1MB of the subtelomere, partitioned by high (≥ 0.85) and low (≤ 0.15) TVR similarity scores (subtelomere proximal 200bp). **g**, Distribution of the mean PCLAI Euclidean Distance in the terminal 1MB of the subtelomere for chromosome arm pairs where the TVR similarity score is greater than or equal to 0.75 at distances of 200, 400 or 600 from the subtelomere junction (* = Mann-Whitney U p < 1 x 10^-10^, underlying swarm plots subsampled to 250 pairs).

To evaluate TVR remodeling in the germline over long timescales, we compared TVR code similarity with local ancestry in the adjacent 1 Mb subtelomeric region using Point-Cloud Local Ancestry Inference (PCLAI) across all 212 HPRC individuals^35^. Telomere arrays with highly similar TVR codes exhibited significantly greater subtelomeric ancestry similarity than telomeres with divergent TVR codes (**Fig. 2f**; median Euclidean distance 0.44, n = 3,201 pairs, versus 2.27, n = 15,624 pairs; Mann Whitney U P = 7.7 × 10^-279^). Furthermore, chromosome ends sharing longer TVR codes exhibited higher subtelomeric ancestry similarity, indicating that the length of TVR similarity reflects the recency of their most recent common ancestral telomere (**Fig. 2g**).

The widespread conservation of subtelomere-proximal TVR codes across the HPRC cohort further demonstrates that these patterns are not sequencing or cell culture artifacts and suggests that telomeres rarely shorten below approximately 200-500 bp in the germline, or that cells harboring shorter telomeres are selectively excluded from transmission. Together, these results indicate that proximal TVR codes form a heritable molecular signature of chromosome ends, whereas distal telomeric regions undergo gradual remodeling during telomere maintenance in the germline.

### Preferential interchromosomal telomere exchange on certain chromosome ends

Rare instances of highly similar TVR codes were observed among telomeres with divergent subtelomeric local ancestry (**Fig. 2f**), indicating that TVR code similarity could uncover recombination events occurring at extreme chromosome termini. Human subtelomeres are known hotspots of interchromosomal recombination and segmental duplication^36^, yet whether the sequence of the telomere array itself is maintained after interchromosomal exchange remains unresolved. Using the proximal 200 bp TVR codes to quantify the extent of interchromosomal end exchange involving telomere arrays revealed that 0.03% of all interchromosomal end pairwise comparisons shared ≥80% similarity (13,762 of 42,925,877 interchromosomal pairs), whereas 0.82% of all intrachromosomal end pairwise comparisons shared ≥80% similarity (9,877 of 1,203,873 intrachromosomal pairs) (**Fig 3a-c; Extended Data Fig. S12a**). These results indicate that chromosome end-specific TVR codes are largely preserved through inheritance from common ancestral chromosome end telomere arrays, which in rare cases derive from different chromosomes via interchromosomal end exchange of telomere arrays.

**Fig. 3.**
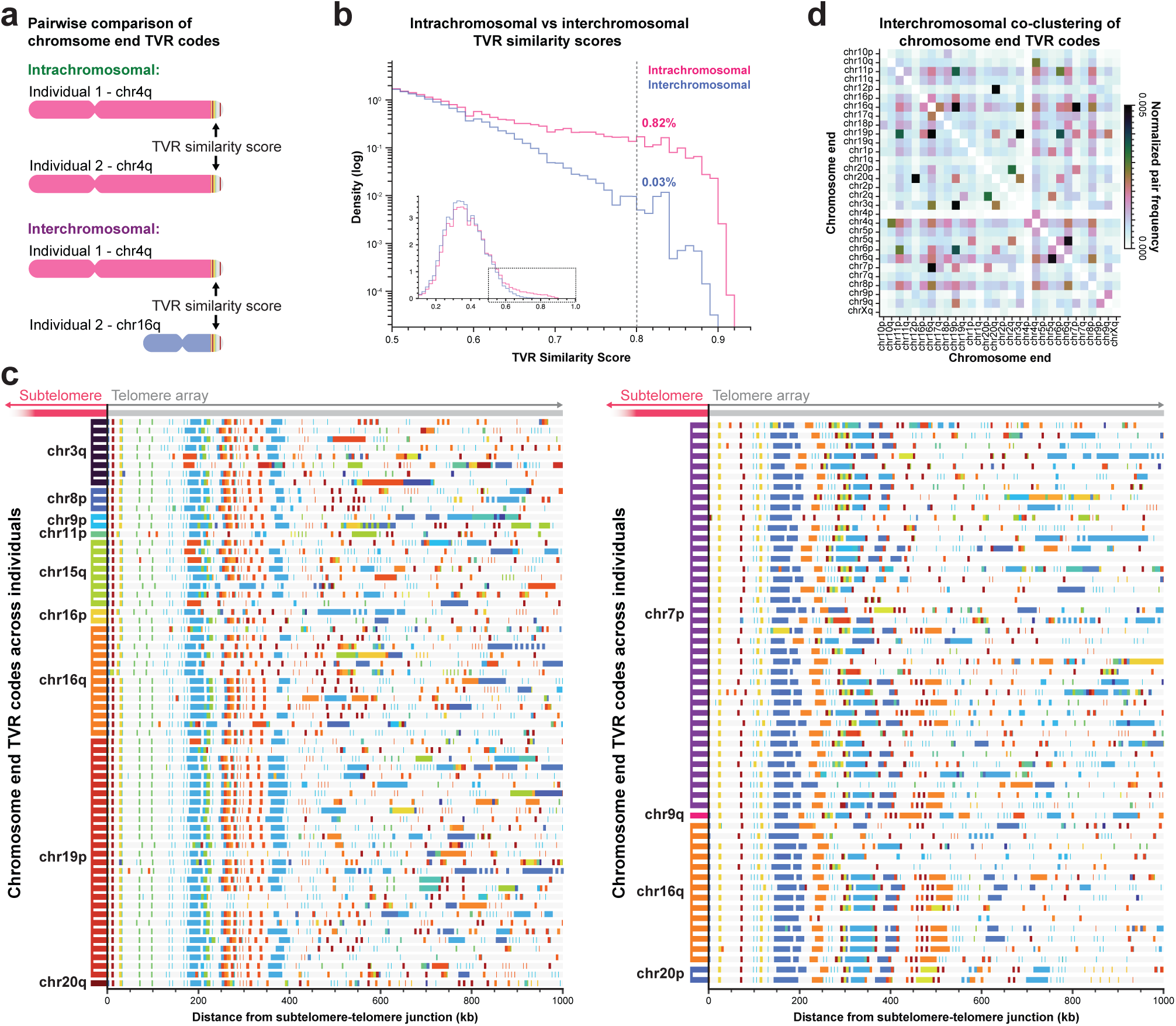
Detection of interchromosomal telomere exchange with TVR codes. **a**, Schematic depicting TVR code comparisons to look for terminal interchromosomal rearrangements. **b**, Distribution of TVR similarity scores (subtelomere proximal 200bp) between chromosome pairs aligned to the same arm (n = 1,203,873 pairs; 9,877 pairs with TVR similarity score ≥ 0.8) and those aligned to different arms (n = 43,925,877 pairs; 13,762 pairs with TVR similarity score ≥ 0.8). **c**, Two examples of TVR codes shared across multiple chromosome arms, each row is a randomly sampled molecule for each haplotype chromosome arm. Minimum TVR similarity score (proximal 200bp) of 0.7 used to generate clusters. **d**, Heatmap of highly similar (score ≥ 0.7) interchromosomal TVR code co-occurrence within the proximal 200 bp. Counts are normalized by the product of the number of haplotypes for both arms (the maximum number of pairs).

In rare cases, shared interchromosomal end TVR codes maintained sequence homology extending up to ∼500 bp into the telomeric repeat (**Fig. 3c; Extended Data Fig. S12a**), consistent with relatively recent interchromosomal end telomere exchanges. Clustering of TVR codes revealed recurrent relationships among chromosome arms 19p and 16q, 7p and 16q, and 20q and 12p, consistent with prior work focused on 16p/16q^11^ (**Fig. 3d**). Alignment of 100 kb of subtelomeric sequence from a subset of chromosome arms with demonstrated shared structural features (**Extended Data Fig. S12b**), suggests that these TVR similarities likely arise from larger subtelomeric interchromosomal rearrangements rather than isolated telomere exchange events. Together, these findings demonstrate that TVR codes provide a sensitive readout of both intra- and interchromosomal end telomere dynamics and that both intrachromosomal and interchromosomal telomere exchange is occurring in the human germline.

### Telomerase-independent telomere lengthening in the human germline

Beyond revealing recombination between chromosome ends, TVR codes may also provide a potential marker for telomerase-independent telomere extension in the human germline. Telomerase-independent telomere maintenance was first described in yeast, where recombination-associated pathways can sustain telomere length in the absence of telomerase^37^. Analogous Break Induced Replication (BIR)-mediated alternative lengthening of telomeres (ALT) mechanisms have since been identified in human cells, particularly in a subset of cancers, where telomere extension occurs through recombination-mediated copying of telomeric DNA using diverse templates, including extrachromosomal telomeric DNA and interchromosomal crossover events^38–42^. Among the 2,408 chromosome-end telomere arrays that met coverage and TVR code length thresholds, 2.08% (50 telomere arrays) exhibited a distinctive pattern of internal TVR duplications consistent with a telomerase-independent extension event (**Fig. 4a; Extended Data S13a**). These events were distributed across chromosome arms, with 23 arms containing at least one such duplication and 37.3% of individual haplotypes harboring at least one affected telomere array. Notably, chromosome arm 19p accounted for 20% of all detected events, and these duplications shared an identical TVR code across unrelated individuals, indicating that this event did not arise from a somatic event during cell culture, but rather, originated in the germline from a common ancestral telomere array (**Fig 4b**). Furthermore, 3 of the 50 telomere arrays harboring an internal TVR duplication showed evidence of nested duplication events (**Extended Data Fig. S13b-d**), wherein a TVR duplication likely occurred twice during that telomere array’s history.

**Fig. 4.**
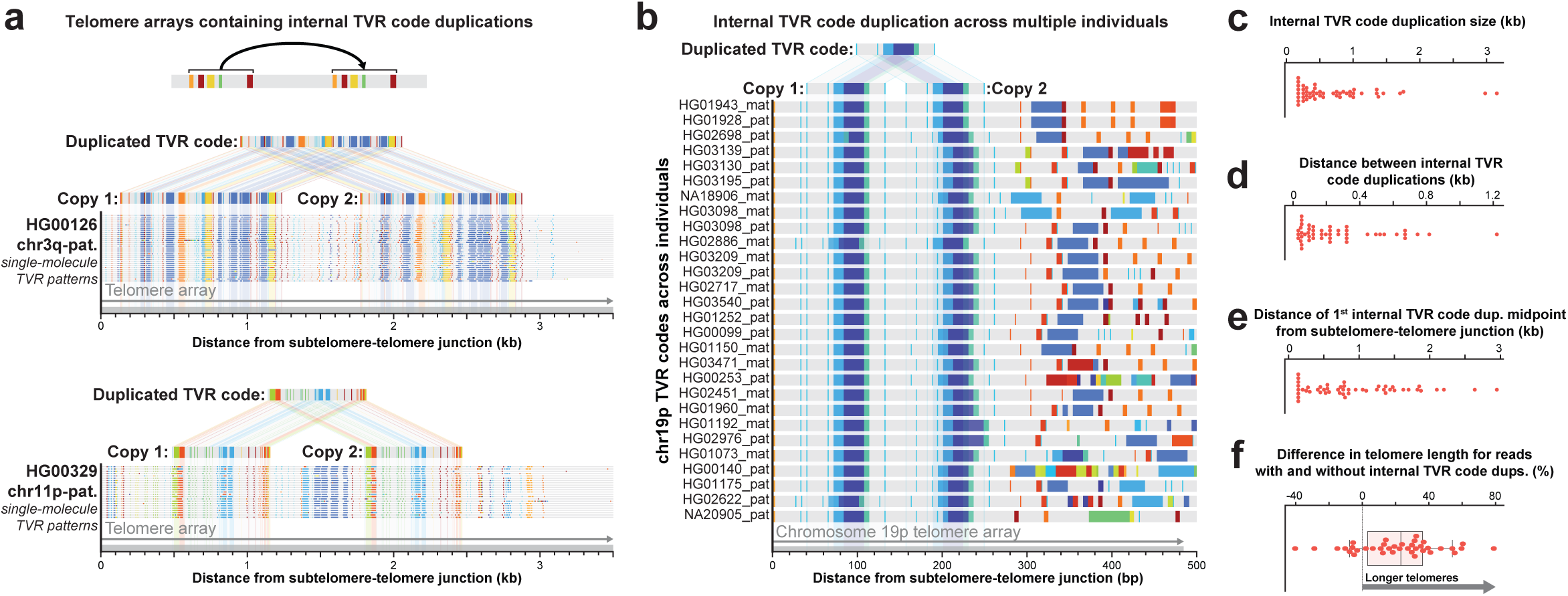
Detecting rare Alternative Lengthening of Telomeres (ALT)-like lengthening in the human germline with TVR codes. **a**, Two examples of internal TVR duplications across molecules in different individuals at different chromosome arm ends (ONT R10). **b**, Example of TVR code duplication event that is present across haplotypes, each haplotype represented by a randomly subsampled molecule from that haplotype chromosome end (PacBio HiFi) **c**, Size of internal TVR duplication event. **d**, Distance between TVR duplication events (start of 2^nd^ internal TVR code - end of 1^st^ internal TVR code). **e**, Distance of 1^st^ internal TVR midpoint to the telomere-subtelomere boundary. **f,** Normalized difference in relative telomere length between chromosome ends with and without internal TVR duplications. Each point represents a haplotype harboring at least one internal TVR duplication. Values indicate the percentage change in median molecule length for arms containing a TVR duplication compared to the median molecule length of all arms without a TVR duplication, within the same haplotype.

Duplicated TVR codes had a mean length of 667 bp and were separated by an average of 239 bp between repeated segments (**Fig. 4c, d**). Notably, these duplications often impacted centrally located TVRs within the telomere array, with a mean midpoint position 1340 bp from the subtelomeric boundary (**Fig. 4e**). Overall, telomere arrays harboring TVR duplication events were 23.12% longer than non-duplicated telomeres from the same individual (**Fig 4f**, median absolute difference 1,259 bp; Wilcoxon signed-rank test, P = 1.74 × 10⁻□), consistent with TVR duplications arising from a mechanism that enables telomere elongation. Together, these findings demonstrate that telomerase-independent telomere extension is operating in the human germline, shaping telomere length variation, structural diversity, and long-term stability across human populations.

### Chromatin compaction of human telomere arrays

As TVRs markedly alter the sequence composition of the proximal telomere array, we next sought to identify whether TVRs adopt a chromatin architecture consistent with that of the telomere cap, or instead enable the encroachment of subtelomeric chromatin into the telomere array. Telomeric DNA is known to be packaged into specialized chromatin in which nucleosome arrays coexist with the Shelterin complex^43^, which binds telomeres in a TTAGGG sequence-specific manner^44,45^. To determine the chromatin architectures adopted by TVRs, we leveraged ONT Fiber-seq data from 22 HPRC samples to co-map telomeric sequence features and protein occupancy events across molecules spanning the telomere-subtelomere boundary (**Fig. 5a**). Using this Fiber-seq dataset, we identified 26,972 molecules spanning the telomere-subtelomere boundary and identified protein occupancy events on each molecule (**Fig. 5a,b**). Subtelomeric regions exhibited regularly spaced mononucleosome arrays consistent with genome-wide Fiber-seq patterns^46^. In contrast, telomeric regions were strongly protected from methylation by the MTase (**Fig. 5c,e-g; Extended Data Fig. S15a,b**), showing an 11.6-fold enrichment relative to subtelomeric DNA for extended MTase-protected footprints exceeding 500 bp (**Extended Data Fig. S15b**).

**Fig. 5.**
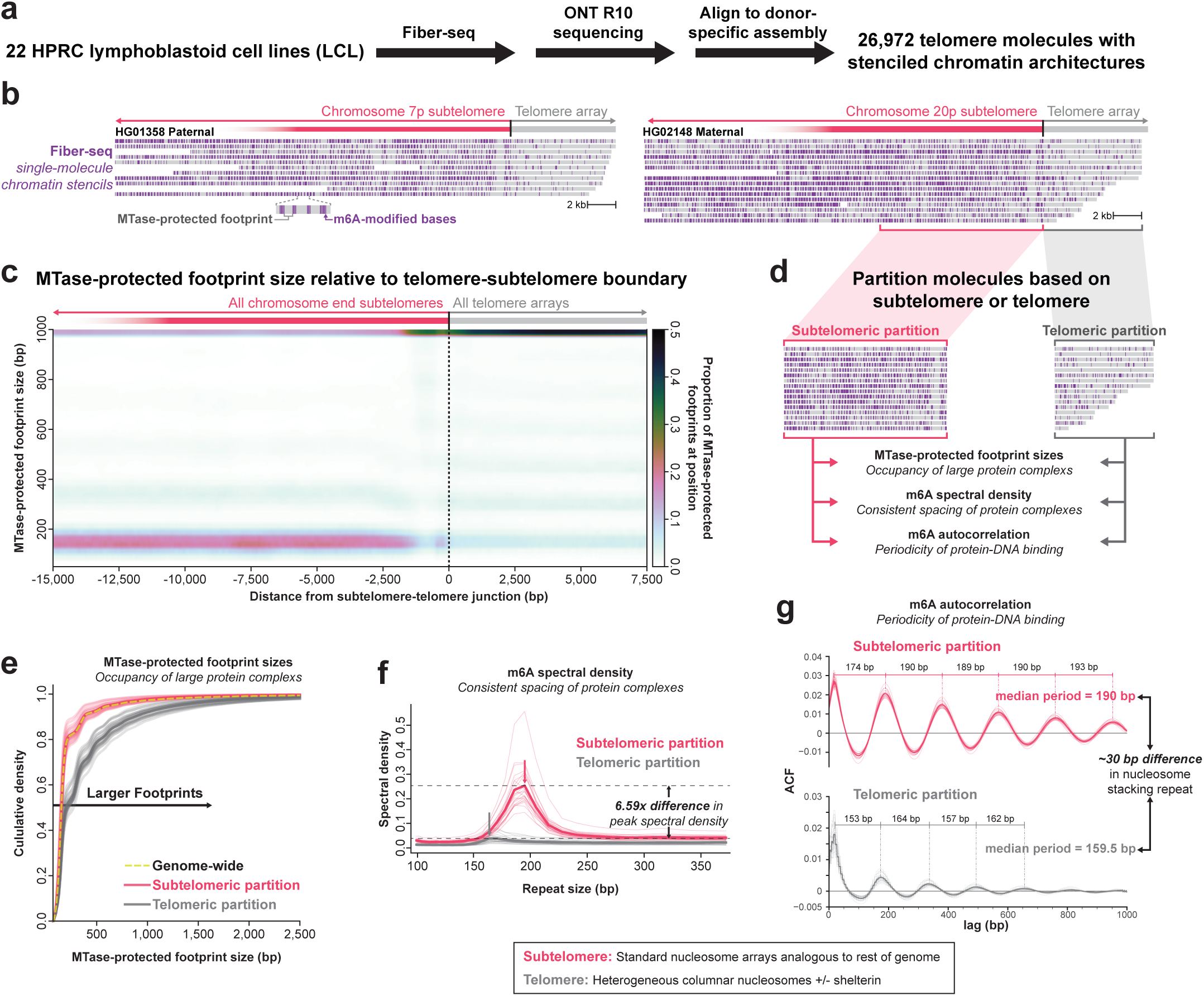
Quantifying telomeric chromatin organization across 22 samples with Fiber-seq. **a**, Overview of Fiber-seq analysis of telomeres in HPRC **b**, Two examples of ONT R10 Fiber-seq single-molecules and m6A methylation events at different chromosome arm termini. Every gray bar is a single-molecule, and every vertical purple line is an m6A event along that molecule. **c**, Heatmap of relative MTase-protected patch size distributions in relation to the subtelomere-telomere junction (in molecular coordinates) across 22 HPRC Fiber-seq samples (mean across all samples shown). MTase-protected patch sizes (y axis) and the positions they cover (x axis) are partitioned in 25-bp and 5-bp bins, respectively. **d**, Schematic describing the splitting of Fiber-seq molecules into their subtelomeric and telomeric components. **e**, Cumulative density of MTase-protected patch sizes. Faint lines represent individual Fiber-seq samples (n = 22), and bold lines indicate the mean distribution across all samples. The genome-wide distribution represents a 1% sample of all molecules genome-wide, filtered to exclude those overlapping repetitive elements. **f,** Spectral density of the m6A signal in subtelomeric and telomeric domains derived from the same molecules (the 4096 bp adjacent to the subtelomere-telomere junction for both the subtelomere and telomere). Faint lines represent individual Fiber-seq samples, and bold lines indicate the mean spectral density across all samples. **g,** Autocorrelation (ACF) of the m6A signal in subtelomeric and telomeric domains. Faint lines represent individual Fiber-seq samples, and bold lines indicate the mean ACF across all samples. ACF smoothed with a 30bp sliding window, and limited to the first 10kb of the telomere.

Despite this extensive protection, 66% of telomere molecules retained at least one mononucleosome-sized footprint within the telomeric repeat (**Extended Data Fig. S15c**), including footprints located near the distal end of the array, consistent with nucleosomes contributing to the telomere cap. To determine whether these large MTase-protected footprints correspond to stacked multimeric nucleosome arrays, we performed autocorrelation and spectral density analyses across telomeric and subtelomeric regions of each molecule^25,30^ (**Fig 5d,f,g**). Compared to adjacent subtelomeres, telomeric regions displayed markedly reduced spectral periodicity, indicating that most extended footprints do not correspond to canonical well-structured nucleosome arrays (**Fig. 5f**). However, when nucleosome multimers were detected within telomeres, they exhibited a distinct nucleosome repeat length (NRL) of ∼159 bp (**Fig 5g**), closely matching the 157 bp NRL previously reported for human telomeric columnar nucleosome stacking^47^, and along rat telomeres, which are an order-of-magnitude longer than human telomeres^48,49^. Together, these results demonstrate that Fiber-seq can resolve both single and multimeric protein occupancy along individual telomeres with single-molecule precision. Furthermore, these results reveal that telomeric chromatin is highly heterogeneous in both footprint size and spatial organization, with the irregular positioning of extended MTase-protected regions suggesting that Shelterin complexes and compacted nucleosome stacks are not uniformly phased along the telomere array.

### TVRs modulate telomere cap chromatin architectures

Having demonstrated that Fiber-seq can accurately measure telomere cap chromatin architecture, we next evaluated whether TVRs adopt a chromatin architecture consistent with that of the telomere cap, or instead enable the encroachment of subtelomeric chromatin into the telomere array. We observed that accessible patches of chromatin between large MTase-protected footprints preferentially localized to the subtelomere-proximal portion of the telomere array (**Fig S15e**, 22.3% increase in the first 2.5 kb vs 2.5-5kb, Wilcoxon signed-rank test, P = 4.77×10^-7^), a region enriched for TVRs. We next quantified the relative chromatin accessibility of TVRs and canonical TTAGGG tracts within telomere arrays, which revealed that the majority of TVRs exhibit significantly higher local chromatin accessibility in the telomere than canonical TTAGGG repeats (**Fig. 6a; Extended Data Fig. S16a,b**), even after controlling for their position within the telomere array (**Fig. 6b**). Notably, TVRs were associated with increased local chromatin accessibility throughout the proximal ∼5 kb of the telomere array, indicating that TVRs locally influence chromatin architectures across a substantial portion of the telomere array (**Fig. 6b**).

**Fig. 6.**
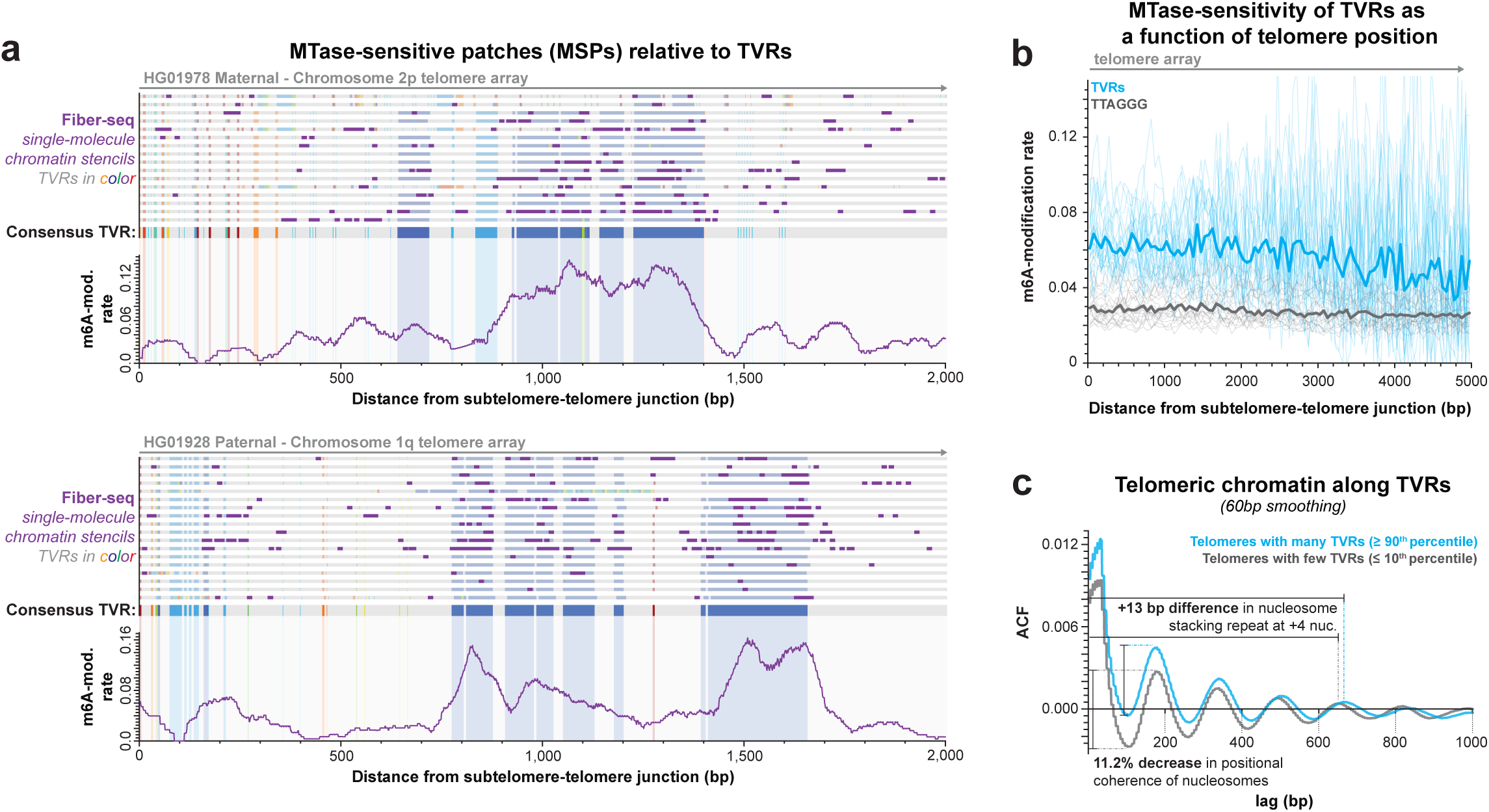
Integration of single-molecule accessibility and sequence information. **a**, Two examples of Fiber-seq m6A deposition along single molecules, overlaid with single-molecule TVR data. The overall m6A/A methylation rate (100 bp smoothing) is shown below each set of molecules. **b,** m6A methylation rate at TVRs versus canonical TTAGGG repeats in 50 bp bins across the telomere. Faint lines represent individual HPRC Fiber-seq samples, and bold lines indicate the cross-sample median. **c**, Autocorrelation analysis of m6A methylation within telomeric segments (60-bp smoothing). Molecules are partitioned into the top and bottom deciles based on their total TVR content.

As telomeres harbor a distinct pattern of nucleosome array stacking as part of the telomere cap (**Fig. 5g**), we next quantified whether TVRs adopt nucleosome array architectures consistent with subtelomeric or telomeric chromatin. Autocorrelation analyses of single-molecule Fiber-seq reads stratified by their overall TVR burden revealed that telomeres with a high TVR load overall exhibited a nucleosome repeat unit that was markedly more consistent with that of telomere columnar stacking as opposed to standard subtelomeric nucleosome arrays (**Fig. 6c**). However, relative to telomere arrays with minimal TVRs, telomere arrays with a high TVR load exhibited a modest attenuation in the peak-to-trough amplitude of the autocorrelation function, as well as a modest increase in the NRL size of a nucleosome multimer (**Fig. 6c**). Together, these findings indicate that although columnar nucleosome stacking does occur overlying TVRs, TVRs nonetheless locally disrupt the continuity of this stacking while largely retaining the configuration. Together, these results demonstrate that TVR-rich subtelomere-proximal regions do not appear to behave as a broadly distinct and separable chromatin domain from the rest of the telomere array. However, by locally increasing DNA accessibility and perturbing nucleosome stacking, TVRs introduce discrete discontinuities into the otherwise compact telomeric chromatin fiber, providing a mechanistic explanation for how underlying telomeric sequence variation can influence the structural organization of chromosome ends.

## Discussion

We find that many human chromosome ends carry a conserved sequence architecture within the proximal telomere, reflected in distinct TVR patterns that are shared across individuals and traceable to ancestral human telomeres. Proximal TVRs represent static features of chromosome ends that can be maintained over long timescales in the germline, providing insights into genomic events molding telomere arrays within the human germline, as well as the impact of telomere sequence variation on telomere cap formation.

Although telomeres undergo cycles of shortening and elongation in the germline, proximal TVR codes remain remarkably stable. This pattern suggests that telomeres rarely erode beyond a proximal region of several hundred base pairs in the germline. These observations provide a potential explanation for the longstanding paradox that telomere length is strongly heritable across human populations despite ongoing telomere turnover^4,31–33^. Rather than being repeatedly rebuilt from a homogeneous template, telomeres appear to maintain a persistent proximal sequence architecture that anchors chromosome-end identity while allowing distal telomeric repeats to undergo gradual remodeling during maintenance. More broadly, these findings raise the possibility that inherited variation in TVR architecture itself may contribute to the heritability of telomere traits.

Our study also highlights an important technical point: current linear reference genomes, including GRCh38, CHM13, and HG002, are poor substrates for accurately studying donor-specific telomeres and subtelomeres, and even high-quality donor-matched diploid assemblies frequently misassemble telomere array sequences. These limitations have important consequences for methods that rely on assigning telomere-containing reads to chromosome ends using a common subtelomere reference^13,14,50,51^. For example, these common reference-based approaches would obscure interchromosomal telomere exchange events. More complete and diverse pangenome representations will probably improve reference-based telomere anchoring, but our results suggest that direct single-molecule read-level sequence evaluation against donor-specific assemblies will remain essential for understanding chromosome-end-specific telomere dynamics.

Using TVR codes as molecular records of telomere history, we identified rare interchromosomal telomere exchange events and recurrent internal duplications within telomere arrays. Importantly, our chromatin data confirmed that these events are occurring within a chromatin architecture consistent with that of the telomere cap. Identical interchromosomal telomere exchange events and internal duplications were present across multiple individuals, indicating that these telomerase-independent telomere lengthening events are operating as part of normal human germline telomere maintenance. These events resemble recombination-associated extension pathways described in yeast and in ALT-positive cancers, but our findings show that these mechanisms can act in humans outside overt pathological contexts. One plausible model is that these duplication events arise through BIR between sister chromatids, analogous to telomere sister-chromatid exchange, facilitated by extensive local homology within telomeric and subtelomeric repeats. Consistent with this model, sister-chromatid exchange and DNA recombination proteins have been observed along telomeres that elongate via a telomerase-independent mechanism in early mouse embryos^52^. Alternatively, these duplication events may reflect replication slippage or template switching within the unusual chromatin environment of chromosome ends. Distinguishing among these mechanisms will require direct observation of telomeric replication and recombination intermediates.

Our results provide new insight into the architecture of the human telomere cap. Single-molecule Fiber-seq profiling shows that telomeric chromatin is heterogeneous and irregularly organized. When nucleosome multimers are present, they adopt a nucleosome repeat length consistent with previously described columnar telomeric nucleosome stacking^47^, providing further experimental evidence in support of this unique telomeric chromatin state. This heterogeneous organization may allow telomeres to satisfy two competing demands: maintaining a protective cap while preserving sufficient structural flexibility for replication and elongation. Importantly, TVRs act as sequence-level modulators of this architecture, introducing localized discontinuities into an otherwise compact telomeric chromatin fiber. In this framework, the TVR code does not simply mark telomere history, but helps shape how shelterin and nucleosomes assemble along telomere arrays, generating reproducible regions of increased chromatin accessibility that may facilitate engagement of the replication machinery. Through this effect on telomere cap chromatin, proximal TVRs may contribute to the faithful maintenance of the proximal telomere and subtelomere, providing a mechanistic basis for the persistence of ancestral TVR codes across the human population.

The observation that TVR abundance is significantly elevated along TAR1-containing chromosome ends suggests a functional coupling between the subtelomere and telomere. Because TAR1 elements encompass the promoter of the TERRA long non-coding RNA, enrichment of proximal TVRs at these chromosome ends may promote the stable maintenance of TAR1-associated subtelomeric genetic architectures. In addition to this, we hypothesize that proximal TVRs directly influence TERRA biology by altering TERRA RNA secondary structure, G-quadruplex stability, and recruitment of TERRA-binding proteins. In this model, the TVR code serves as a sequence-encoded interface between subtelomeric transcription and telomere cap architecture. Future studies leveraging the atlas of TVR codes presented here will enable direct tests of how specific TVR sequences influence TERRA structure and function.

Taken together, our results show that human telomeres contain a sequence-encoded architecture that links chromosome-end sequence variation to telomere chromatin organization and inheritance across the human population. By integrating long-read sequencing with single-molecule chromatin profiling across hundreds of matched donor-specific assemblies, this study provides a framework for understanding how telomere sequence variation contributes to chromosome-end biology and establishes new principles governing the maintenance and long-term persistence of human telomeres.

## Methods

### Code availability

All code to recreate analysis and figures in this paper is available at: https://github.com/StergachisLab/HPRC-R2-Single-Molecule-Telomere-Analysis.

### Data Availability

All HPRC data, including assemblies, PacBio HiFi reads and R10 ONT reads are available at: https://data.humanpangenome.org/raw-sequencing-data.

Fiber-seq data is available at https://s3-us-west-2.amazonaws.com/human-pangenomics/index.html?prefix=submissions/5ECA1D3E-1C37-44B7-BA20-576417F786F0--UCSC_HPRC_ONT_YEAR1_FIBERSEQ/ .

Processed files are available at: https://github.com/StergachisLab/HPRC-R2-Single-Molecule-Telomere-Analysis.

### Filtering for HPRC telomeric reads

We acquired all PacBio HiFi and Oxford Nanopore Technologies (ONT) R10 sequencing data associated with the Human Pangenome Reference Consortium (HPRC) samples, alongside their corresponding assemblies (available at: https://github.com/human-pangenomics/hprc_intermediate_assembly/tree/main/data_tables). ONT R9 data was excluded from this analysis due to excessive sequencing errors observed in the telomeric regions (unpublished). To define our telomeric boundaries, we identified terminal telomeric repeat stretches using seqtk^1^ telo (v1.4-r122). We extracted reads overlapping telomeric regions using samtools^2^ (v1.18), applying the flags -F 2308 -L <SEQTK telo output< directly to the AWS S3 BAM URLs. We then isolated high-quality telomeric reads by requiring a mean PHRED score greater than 20 along the query coordinates extending past the telomeric boundary. To assign a reference chromosome designation to each contig we utilized the curated chromosome assignment files from HPRC : https://github.com/human-pangenomics/hprc_intermediate_assembly/tree/main/data_tables/annotation. Finally, to orient the reads in the proper direction and embed them into a shared molecular coordinate space, we assigned all p-end telomeres (-) directionality and q-end telomeres (+) directionality and then centered the filtered BAM files at the telomere boundary using fibertools^3^ (ft) center with the parameters -t 30 -b <SEQTK telo output> -w -F 2308. From this ft center output, we extracted telomeric sequences across all individuals, ensuring all molecular coordinates were anchored at their respective subtelomere–telomere boundary (represented as position 0 in the telomeric array).

### Benchmarking alignments in CHM13, HG002, and with a donor specific assembly

To quantify the benefit in a donor-specific assembly (DSA) compared to standard telomere-resolved references, we extracted reads that mapped to the terminal 100 kb of HG00099. These reads were then realigned to both the CHM13 (chm13v2.0) and HG002 (hg002v1.0.1) reference assemblies using minimap2^4^ (-k 15 -a -x map-ont --cs --eqx -L). To visualize sequencing depth at the chromosome termini, we generated bigWig tracks using samtools depth -Q 30 followed by bedGraphToBigWig and visualized the tracks in IGV^5^. The quality of the resulting primary alignments was quantified by filtering for minimum mapping quality (MAPQ) thresholds of 20, 30, 40, 50, and 60 using samtools view -c -F 2308 -q <CUTOFF>. Finally, to assess the impact of telomeric sequence on mapping fidelity across all Fiber-seq samples, we grouped alignments into telomere-overlapping and non-telomere-overlapping groups, counting the number of mismatches (NM tag in the BAMs) per group.

### Quantifying amount and distribution of telomere boundary spanning reads

Alternative or unplaced contig assignments were removed using regular expression matching. Each read was then assigned to a specific chromosome arm (p or q) based on a genomic coordinate threshold of 100 kb relative to the centering position (p < 100kb telomere start, q > 100kb telomere start).

### Identifying TVRs

We define Telomere Variant Repeats (TVRs) as any sequence that impedes the continuous pattern of canonical TTAGGG placement along the telomere. To identify TVRs along single molecules, we extract the telomeric portion of every read and then apply a simple regular expression for the canonical TTAGGG repeat and flag any gaps where this motif is absent. For robust identification of specific variant repeat types, please refer to Grigorev et al. (2021)^6^ and Stephens et al. (2024)^7^.

### Visualization and color mapping of telomeric variant repeats

To ensure consistent visual representation of TVRs across single-molecule plots, we implemented a deterministic, hash-based color mapping algorithm. For each mapped 6-mer, the sequence is evaluated alongside its reverse complement, and the lexicographically smaller sequence is selected as a representative key to guarantee strand-agnostic color assignment. To accommodate variants of varying lengths, TVRs longer than 6 bp are partitioned into consecutive 6-bp windows (window size = 6bp, and step size = 6bp) for color assignment, while TVRs shorter or equal to 6 bp in length are colored based on their entire sequence. This representative key is then hashed into an integer and the resulting integer is normalized to a scalar value between 0 and 1, which is subsequently mapped to a continuous color palette (the matplotlib ‘turbo’ colormap).

### Quantifying TVR abundance, location, and composition

Inspection of TVRs greater than 300bp showed many molecule-specific homopolymer stretches, likely corresponding sequencing artifacts, so we limited our analysis of TVRs to those from 1-300bp. Because every molecule was anchored at the subtelomere-telomere junction we calculated the absolute number of reads at every position (variable due to different molecular telomere lengths) and then the absolute count of bases that are classified as a TVR at every position, and we can then calculate the TVR depth divided by absolute depth to get the TVR composition per position relative to the boundary. To compare the 6-mer composition of telomere variant repeats (TVRs) across chromosome ends, we extracted TVR coordinates (minimum TVR size of 6bp) from every molecule assigned to a specific chromosome end, limited to TVRs located within the proximal 5 kb of the telomeric array. We partitioned each TVR into non-overlapping chunks of size 6bp. For each chromosome arm, we aggregated the total number of base pairs corresponding to each unique 6-mer sequence and, to enable comparisons between arms, normalize these sums by the total number of sequencing reads for that chromosome arm. For plotting we calculated the global abundance of all 6-mers across all chromosome ends and kept the top 10 most frequent 6-mers, aggregating all other 6-mers into a single combined category..

### TAR1 stratified analysis

We downloaded repeat annotations for each assembly from https://github.com/human-pangenomics/hprc_intermediate_assembly/blob/main/data_tables/annotation/repeat_masker/repeat_masker_bed_hprc_r2_v1.0.index.csv and extracted all TAR1 containing annotations with a grep search for “TAR1”. We then assigned each chromosome end as TAR1+ or TAR1-based on whether there was a TAR1 element within 10kb of the telomere/subtelomere boundary.

### Fiber-seq processing (CHM13 / HG002)

CHM13 and HG002 Fiber-seq data were downloaded from GSM7074431 and GSM7074433, respectively^8^. Data was reprocessed (ft call-m6a, ft add-nucleosomes) as suggested in Jha and Bohaczuk et. al. 2024^3^. Reads were subsequently anchored at telomere boundary with ft center as described above.

### CTCF analysis

To identify CTCF motifs, we utilized FIMO^9^ with the JASPAR^10^ position weight matrix for MA0139.1 applying the following parameters: --no-qvalue, --thresh 0.001, and --max-stored-scores 1000000. For the HPRC assemblies, downstream analysis was strictly limited to motifs with a score of 60 or greater. We restricted our CTCF footprinting analysis to the CHM13 line with PacBio HiFi sequencing because PacBio HiFi sequences both strands, both adenine (A) and thymine (T) positions are informative, whereas single-stranded ONT only captures modifications on A bases. For footprinting in CHM13, we required a minimum coverage of 10 overlapping molecules per CTCF motif. We then executed fibertools footprint, utilizing a YAML configuration file that defined the entire motif as a single continuous module ([0,19]).

### Pedigree analysis

To evaluate the heritability of TVR codes, we analyzed long-read HiFi sequencing data from the Genome in a Bottle^11^ (GIAB) trio (HG002: son; HG003: father; HG004: mother). Alignments to hg38 categorized under “PacBio CCS 15–20 kb Chemistry v2.0” were retrieved from the GIAB data index repository (https://github.com/genome-in-a-bottle/giab_data_indexes). We first extracted all reads mapping to the terminal 50 kb of each chromosome. All extracted reads were realigned to the HG002 reference assembly (hg002v1.0.1) using minimap2 (-ax map-hifi --MD -c-y). We then filtered the data to retain only primary alignments (samtools view -b -F 2308) and centered these alignments at the telomere boundary using ft center. Finally, TVRs were plotted as described above with the exception that the minimum TVR size threshold was set to 6 bp to account for sequencing artifacts introduced in telomeric regions by older PacBio chemistries.

### TVR code alignment and clustering

To generate TVR consensus sequences, we first filtered for chromosome ends supported by at least five molecules mapping into the telomere array. We constructed a 2D matrix where each row represents a molecular sequence corresponding to the telomere array, and each column denotes the distance from the subtelomere-telomere boundary (position 0). At each position, the base present in the majority of reads was selected as the consensus base. Finally, we truncated the resulting consensus telomeric sequence to the median length across all constituent molecules. Because this approach lacks an inter-molecule alignment step, the quality of the consensus sequence progressively degrades moving distally into the telomere.

To align consensus telomere sequences based on their TVR position and content, we implemented a custom local alignment algorithm that specifically ignores standard canonical sequence. First, we slide a window across each consensus sequence and extract overlapping segments of 6 bases (k=6). To force the alignment to utilize TVRs, we identify all canonical telomeric hexamers (exact matches to TTAGGG or CCCTAA). During the alignment, if a 6-mer in either sequence belongs to this canonical set, it is completely masked and contributes a score of 0. For the remaining non-canonical sequence, we use a Smith-Waterman^12^ dynamic programming approach to calculate alignment scores. We award a score of 2.0 for a match, while penalizing mismatches by -1.0 and gaps by -2.0 between TVRs. Finally, to compare similarities across variable length stretches of telomere sequence (200bp from subtelomere-telomere junction, 400bp, etc.), we then normalize the raw alignment score at specific truncation lengths. We do this by dividing the maximum observed score by the theoretical ideal score (the match score of 2.0/bp at TVRs multiplied by the number of TVR 6-mers in the shorter sequence), producing a final similarity metric that scales from 0 to 1.

To group telomeres based on TVR code similarity across distinct haplotypes, we hierarchically clustered the haplotype-specific chromosome ends based on their pairwise similarity scores calculated at each specific sequence truncation length (200bp from subtelomere-telomere junction, 500bp from subtelomere-telomere junction, etc.). For each chromosome arm, a similarity matrix was constructed from the pairwise alignment data, with self-similarity along the diagonal fixed to 1. This similarity matrix converted into distance matrix, where distance was defined as 1 minus the similarity score. We then applied the Unweighted Pair Group Method with Arithmetic Mean (UPGMA^13^) hierarchical clustering algorithm to the distance matrix (implemented via scipy.cluster.hierarchy.linkage with method=’average’). Finally, the optimal one-dimensional order of the resulting dendrogram leaves (scipy.cluster.hierarchy.leaves_list) was extracted to group structurally similar telomeric alleles adjacent to one another for visualization. To perform interchromosomal clustering, we used the same workflow described above.

### Local ancestry analysis with Point Cloud Local Ancestry Inference (PCLAI)

We utilized Point Cloud Local Ancestry Inference (PCLAI)^14^ to characterize the local ancestry at the terminal subtelomere. Briefly, PCLAI embeds ancestry information (projected in CHM13 reference space) into a principal component (PC) space, where genomic windows encompassing 1,000 SNPs are represented by their top 2 principal components (PC1 and PC2). We restricted our focus to windows situated entirely within the terminal 1 Mb of each subtelomere. To quantify the genetic distance between any two subtelomeric regions, we first calculated the Euclidean distance between bins across this 1 Mb window for each haplotype-specific chromosome arm. We then used these distance values to compute the mean Euclidean distance between all bins in the subtelomeric 1MB, providing a single value for ancestry divergence between pairs of subtelomeres belonging to different haplotypes.

### Alignment of 100kb subtelomeric region

We selected a subset of chromosome ends sharing proximal TVR codes across different chromosome arms for an all-vs-all alignment. For each chromosome end, a 100 kb genomic window extending from the subtelomere-telomere boundary into the adjacent subtelomeric region was used to extract the corresponding nucleotide sequences with samtools faidx and concatenated into a single multi-haplotype FASTA file. An all-vs-all pairwise sequence alignment of the concatenated sequences was performed using minimap2 with the parameters - t 32 -x asm20 -c -eqx -D -P --dual=no. The resulting alignments were subsequently visualized using SVbyEye^15^.

### Alternative lengthening mechanism quantification description

Traditional methods for identifying structural duplications are ineffective for telomere variant repeats (TVRs) because the TTAGGG sequences flanking these units are also homologous. To address this limitation, we did a manual inspection of 2,408 Oxford Nanopore Technologies (ONT) R10 telomere arrays to identify instances of TVR duplications. Two independent reviewers manually annotated these repeat instances. To establish consensus boundaries, we calculated the mean genomic coordinates from the two sets of independent annotations and tabulated the results.

### Fiber-seq processing (HPRC, ONT R10)

Basecalling on R10 data was performed using dorado with basecall_model=dna_r10.4.1_e8.2_400bps_sup@v5.2.0 and modbase_models=dna_r10.4.1_e8.2_400bps_sup@v5.2.0_5mCG_5hmCG@v2,dna_r10.4.1_e 8.2_400bps_sup@v5.2.0_6mA@v1. Reads were then subsequently aligned to their respective assembly using minimap2(v2.29-r1283) -t 32 --secondary=no -I 8G --eqx --MD -Y -y -ax lr:hq. We then filter out low-quality base modifications (the bottom 10%) with modkit call-mods (v0.5) - p 0.1. We identify methyltransferase (MTase) protected patches using ft add-nucleosomes. We then orient and anchor all molecules to their telomere/subtelomere boundary as before with ft center with the parameters -t 30 -b <SEQTK telo output> -w -F 2308. Finally, to exclude reads originating from cells with unsuccessful Hia5 treatment, we removed molecules with an overall m6A density below 0.1% (equivalent to 1 m6A per 1 kb).

### HPRC Fiber-seq chromatin analysis

To analyze the distribution of MTase protected regions across the telomere boundary, single-molecule coordinate data for each HPRC Fiber-seq sample was filtered to retain high-quality molecules, requiring a minimum read length of 15 kb, at least 1.5 kb of telomeric sequence, a minimum of 100 m6A modifications and at least 10 MTase protected regions (putative nucleosomes). Valid protected fragments ranging from 50 to 1,000 bp in length that mapped within a 30 kb window spanning the boundary (−20 kb subtelomeric to +10 kb telomeric) were discretized into a two-dimensional matrix using 5 bp coordinate bins and 25 bp MTase protected patch size bins. We then performed column-normalization to yield the proportional frequency of protected region sizes at each specific coordinate bin. Finally, to establish a generalized cross-sample profile, an aggregate consensus heatmap was generated by calculating the mean normalized proportional frequency across all individual HPRC sample heatmaps at every positional bin. To calculate the spectral density of the telomeric or subtelomeric component along single molecules we iterate through every individual cell line’s set of reads and then filter for reads where both the telomeric and subtelomeric stretch of molecular sequence is greater or equal to 4096bp in length. We select the terminal 4096bp of the subtelomere directly adjacent to the boundary as the subtelomeric segment and the 4096bp telomeric fragment adjacent to the boundary as the segment representative of the telomere for each molecule. We then generate binary arrays for the subtelomeric and telomeric segments where all bases are 0, except those that represent an m6A, which we label as a 1. We then apply the periodogram function from scipy.signal to calculate power of the signal at different base pair frequencies. Similarly, we split reads into their subtelomeric (terminal 10kb proximal to the subtelomere-telomere junction)and telomeric (limited to those greater than 1kb in length and less than 20kb in length) components to calculate the autocorrelation function (ACF) representing periodic spacing of nucleosomes, we calculated the ACF with a maximum lag of 1,000 bp for all reads per sample, and then calculated the mean ACF across individuals.

### TVR Methylation Rate

To compare the overall distribution of Fiber-seq methylation in TVRs against canonical TTAGGG repeats, we calculated per-molecule measurements of TVR and canonical methylation rates. For each molecule extending into the telomere, the methylation rate was computed by dividing the number of observed methylation events by the total number of potential target bases (ONT data reads out a single strand, so methylations are on A or T dependent on which strand passes through the pore) within the TVR or canonical segments of that molecule’s telomere array. To account for position-dependent effects, the distance from the subtelomere-telomere junction was partitioned into 50 bp non-overlapping bins. Methylation rates for TVR and canonical TTAGGG sequences were calculated independently for the corresponding sequences within each bin. To generate the ACF for varying TVR burdens, we stratified the molecules by TVR burden, extracted the top and bottom deciles, and performed auto-correlative analysis on these subsets of molecules as described in the previous section. To validate that elevated TVR methylation reflects true chromatin accessibility rather than sequence-specific enzymatic biases, we established a background genomic methylation rate for common TVR 6-mers using a random 5% sample of reads aligning to the entire genome. We then calculated the fold-change between the telomeric methylation rate and this genomic baseline for both TVRs and canonical repeats. The relative enrichment rate was then calculated as this TVR fold-change divided by the canonical fold-change, so that this metric represents the change in accessibility for TVRs at the telomere, properly normalized to the baseline genomic and telomeric behavior of canonical TTAGGG repeats.

## Author contributions

D.D., A.E.S.C., and T.R. performed computational analysis. A.B.S., J.R. and B.M. designed and performed Fiber-seq experiments. D.D., M.R.V. and S.J.N. processed Fiber-seq data.

M.S.D.R.P. and D.D. identified all TVR duplication events. J.K.L contributed to identification and procurement of appropriate HPRC datasets. D.D., N.A. and A.B.S. wrote the manuscript.

## Declaration of Generative AI and AI-assisted Technologies in the Writing Process

During the preparation of this work, the authors used LLMs in order to assist with specific programming tasks, and on rare occasions to improve the language of the manuscript. After using these services, the authors reviewed and edited the content as needed and take full responsibility for the content of the published article.

## Human Pangenome Reference Consortium Version 2 Authors

Derek Albracht^1^, Ivan A. Alexandrov^2^, Jamie Allen^3^, Alawi A. Alsheikh-Ali^4^, Nicolas Altemose^5^, Casey Andrews^6^, Dmitry Antipov^7^, Lucinda Antonacci-Fulton^1^, Mobin Asri^8^, Marcelo Ayllon^9^, Jennifer R. Balacco^10^, Floris P. Barthel^11^, Edward A. Belter Jr^1^, Halle D. Bender^8^, Andrew P. Blair^8^, Davide Bolognini^12^, Katherine E. Bonini^13^, Christina Boucher^14^, Guillaume Bourque^15,16,17^, Silvia Buonaiuto^18^, Shuo Cao^18^, Andrew Carroll^19^, Ann M. Mc Cartney^8^, Monika Cechova^8^, Mark J.P. Chaisson^20^, Pi-Chuan Chang^19^, Xian Chang^8^, Jitender Cheema^3^, Haoyu Cheng^21^, Claudio Ciofi^22^, Hiram Clawson^8^, Sarah Cody^1^, Vincenza Colonna^18^, Holland C. Conwell^23^, Robert Cook-Deegan^24^, Mark Diekhans^8^, Maria Angela Diroma^22^, Daniel Doerr^25,26,27^, Zheng Dong^6^, Danilo Dubocanin^5^, Richard Durbin^28,29^, Jana Ebler^25,30^, Evan E. Eichler^9,31^, Jordan M. Eizenga^8^, Parsa Eskandar^8^, Eddie Ferro^14^, Anna-Sophie Fiston-Lavier^32,33^, Sarah M. Ford^23^, Willard W. Ford^34^, Giulio Formenti^10^, Adam Frankish^3^, Mallory A. Freeberg^3^, Qichen Fu^6^, Stephanie M. Fullerton^35^, Robert S. Fulton^1^, Yan Gao^36^, Gage H. Garcia^9^, Obed A. Garcia^37^, Joshua M.V. Gardner^8^, Shilpa Garg^38^, Erik Garrison^18^, Nanibaa’ A. Garrison^39,40,41^, John E. Garza^1^, Margarita Geleta^42^, Mohammadmersad Ghorbani^43^, Tina A. Graves-Lindsay^1^, Richard E. Green^23^, Cristian Groza^44^, Bida Gu^20^, Andrea Guarracino^11,18^, Melissa Gymrek^45^, Maximilian Haeussler^8^, Leanne Haggerty^3^, Ira M. Hall^46,47^, Nancy F. Hansen^7^, Yue Hao^11^, Mohammad Amiruddin Hashmi^4^, David Haussler^8^, Prajna Hebbar^8^, Peter Heringer^25,26,27^, Glenn Hickey^8^, Todd L. Hillaker^8^, S. Nakib Hossain^3^, Neng Huang^36,48^, Sarah E. Hunt^3^, Toby Hunt^3^, Alexander G. Ioannidis^5,8^, Nafiseh Jafarzadeh^8^, Nivesh Jain^10^, Erich D. Jarvis^10,31^, Maryam Jehangir^11^, Juan Jiang^6^, Eimear E. Kenny^13^, Juhyun Kim^7^, Bonhwang Koo^10^, Sergey Koren^7^, Milinn Kremitzki^1,6^, Charles H. Langley^49^, Ben Langmead^50^, Heather A. Lawson^6^, Daofeng Li^6^, Heng Li^36,48^, Wen-Wei Liao^46,47^, Jiadong Lin^9^, Tianjie Liu^6^, Glennis A. Logsdon^51^, Ryan Lorig-Roach^8^, Jonathan LoTempio Jr^52^, Hailey Loucks^8^, Jane E. Loveland^3^, Jianguo Lu^53^, Shuangjia Lu^46,47^, Julian K. Lucas^8^, Walfred Ma^20^, Juan F. Macias-Velasco^1,6,54^, Kateryna D. Makova^55^, Maximillian G. Marin^36,48^, Christopher Markovic^1^, Tobias Marschall^25,30^, Franco L. Marsico^18^, Fergal J. Martin^3^, Mira Mastoras^8^, Capucine Mayoud^32^, Brandy McNulty^8^, Jack A. Medico^10^, Julian M. Menendez^8^, Karen H. Miga^8^, Anna Minkina^56^, Matthew W. Mitchell^57^, Saswat K. Mohanty^58^, Younes Mokrab^43,59,60^, Jean Monlong^61^, Shabir Moosa^43^, Avelina Moreno-Ochando^62,63^, Shinichi Morishita^64^, Jonathan M. Mudge^3^, Katherine M. Munson^9^, Njagi Mwaniki^65^, Nasna Nassir^4^, Chiara Natali^22^, Shloka Negi^8^, Lingbin Ni^9^, Adam M. Novak^8^, Faith Okamoto^8^, Pilar N. Ossorio^66^, Chie Owa^64^, Sadye Paez^10^, Benedict Paten^8^, Clelia Peano^12,67^, Adam M. Phillippy^7^, Brandon D. Pickett^7^, Laura Pignata^18^, Nadia Pisanti^65^, David Porubsky^9,68^, Pjotr Prins^18^, Anandi Radhakrishnan^8^, T. Rhyker Ranallo-Benavidez^11^, Brian J. Raney^8^, Mikko Rautiainen^69^, Alessandro Raveane^12^, Luyao Ren^9,31^, Arang Rhie^7^, Fedor Ryabov^70,71^, Samuel Sacco^23^, Farnaz Salehi^18^, Michael C. Schatz^50,72^, Laura B. Scheinfeldt^73^, Aarushi Sehgal^34^, William E. Seligmann^23^, Mahsa Shabani^74^, Kishwar Shafin^19^, Shadi Shahatit^32^, Ruhollah Shemirani^13^, Vikram S. Shivakumar^50^, Swati Sinha^3^, Jouni Sirén^8^, Linnéa Smeds^58^, Steven J. Solar^7^, Marco Sollitto^10,22^, Nicole Soranzo^12,28,75^, Andrew B. Stergachis^9,56^, Marie-Marthe Suner^3^, Yoshihiko Suzuki^64^, Arda Söylev^25,30^, Ahmad Abou Tayoun^76,77^, Jack A.S. Tierney^3^, Chad Tomlinson^1^, Francesca Floriana Tricomi^3^, Mohammed Uddin^4,78^, Matteo Tommaso Ungaro^23,79^, Rahul Varki^14^, Flavia Villani^18^, Ivo Violich^8^, Mitchell R. Vollger,^56^, Brian P. Walenz^7^, Charles Wang^80^, Lisa E. Wang^13^, Ting Wang^1,6,54^, Aaron M. Wenger^81^, Conor V. Whelan^10^, Zilan Xin^6^, Zheng Xu^6^, Kai Ye^82^, DongAhn Yoo^9^, Wenjin Zhang^6^, Ying Zhou^36^, Xiaoyu Zhuo^6^, Giulia Zunino^12^

## Human Pangenome Reference Consortium Version 2 Author Affiliations

^1^ McDonnell Genome Institute, Washington University School of Medicine, St. Louis, MO 63108, USA

^2^ Department of Human Molecular Genetics and Biochemistry, Faculty of Medical and Health Sciences, Tel Aviv University, Tel Aviv 69978, Israel

^3^ European Molecular Biology Laboratory, European Bioinformatics Institute (EMBL-EBI), Wellcome Genome Campus, Hinxton, Cambridge CB10 1SD, UK

^4^ Center for Applied and Translational Genomics (CATG), Mohammed Bin Rashid University of Medicine and Health Sciences, Dubai Health, Dubai, UAE

^5^ Department of Genetics, Stanford University, Palo Alto, CA 94304 USA

^6^ Department of Genetics, Washington University School of Medicine, St. Louis, MO 63110, USA

^7^ Genome Informatics Section, Center for Genomics and Data Science Research, National Human Genome Research Institute, National Institutes of Health, Bethesda, MD 20892, USA

^8^ UC Santa Cruz Genomics Institute, University of California, Santa Cruz, CA 95060, USA

^9^ Department of Genome Sciences, University of Washington School of Medicine, Seattle, WA 98195, USA

^10^ The Vertebrate Genome Laboratory, The Rockefeller University, New York, NY 10065, USA

^11^ Bioinnovation and Genome Sciences, The Translational Genomics Research Institute (TGen), Phoenix, AZ 85004, USA

^12^ Human Technopole, Milan, Italy

^13^ Institute for Genomic Health, Icahn School of Medicine at Mount Sinai, New York, NY 10029, USA

^14^ Department of Computer and Information Science and Engineering, University of Florida, Gainesville, FL 32611, USA

^15^ Canadian Center for Computational Genomics, McGill University, Montréal, QC H3A 0G1, Canada

^16^ Department of Human Genetics, McGill University, Montréal, QC H3A 0G1, Canada

^17^ Victor Phillip Dahdaleh Institute of Genomic Medicine, Montréal, QC H3A 0G1, Canada

^18^ Department of Genetics, Genomics and Informatics, University of Tennessee Health Science Center, Memphis, TN 38163, USA

^19^ Google LLC, Mountain View, CA 94043, USA

^20^ Quantitative and Computational Biology, University of Southern California, Los Angeles, CA 90089, USA

^21^ Department of Biomedical Informatics and Data Science, Yale School of Medicine, New Haven, CT 06510, USA

^22^ Department of Biology, University of Florence, Sesto Fiorentino, FI 50019, Italy

^23^ Department of Ecology and Evolutionary Biology, University of California, Santa Cruz, CA 95060, USA

^24^ Arizona State University, Consortium for Science, Policy & Outcomes, Washington, DC 20006, USA

^25^ Center for Digital Medicine, Heinrich Heine University Düsseldorf, Düsseldorf, NRW, DE

^26^ Department for Endocrinology and Diabetology at the Medical Faculty and University Hospital Düsseldorf, Heinrich Heine University Düsseldorf, Düsseldorf, NRW, DE

^27^ Paul-Langerhans-Group Computational Diabetology, German Diabetes Center (DDZ) and Leibniz Institute for Diabetes Research, Düsseldorf, NRW, DE

^28^ Wellcome Sanger Institute, Genome Campus, Hinxton, CB10 1RQ, UK

^29^ Department of Genetics, University of Cambridge, Cambridge, CB2 3EH, UK

^30^ Institute for Medical Biometry and Bioinformatics, Medical Faculty and University Hospital DulJsseldorf, Heinrich Heine University, DulJsseldorf, NRW, DE

^31^ Howard Hughes Medical Institute, Chevy Chase, MD 20815, USA

^32^ ISEM, Univ Montpellier, CNRS, IRD, Montpellier, FR

^33^ Institut Universitaire de France, Paris, FR

^34^ Department of Computer Science and Engineering, University of California San Diego, La Jolla, CA 92093, USA

^35^ Department of Bioethics & Humanities, University of Washington School of Medicine, Seattle, WA 98195, USA

^36^ Department of Data Science, Dana-Farber Cancer Institute, Boston, MA 02215, USA

^37^ Department of Anthropology, University of Kansas, Lawrence, KS 66045, USA

^38^ School of Health Sciences, University of Manchester, Manchester M13 9PL, UK

^39^ Traditional, ancestral and unceded territory of the Gabrielino/Tongva peoples, Institute for Society & Genetics, University of California, Los Angeles, Los Angeles, CA 90095, USA

^40^ Traditional, ancestral and unceded territory of the Gabrielino/Tongva peoples, Institute for Precision Health, David Geffen School of Medicine, University of California, Los Angeles, Los Angeles, CA 90095, USA

^41^ Traditional, ancestral and unceded territory of the Gabrielino/Tongva peoples, Division of General Internal Medicine & Health Services Research, David Geffen School of Medicine, University of California, Los Angeles, Los Angeles, CA 90095, USA

^42^ Department of Electrical Engineering and Computer Science, University of California, Berkeley, Berkeley, CA 94720, USA

^43^ Medical and Population Genomics Lab, Sidra Medicine, Doha, Qatar

^44^ Montreal Heart Institute, Montréal, QC, Canada

^45^ Department of Pediatrics, University of California San Diego, La Jolla, CA 92093, USA

^46^ Center for Genomic Health, Yale University School of Medicine, New Haven, CT 06510, USA

^47^ Department of Genetics, Yale University School of Medicine, New Haven, CT 06510, USA

^48^ Department of Biomedical Informatics, Harvard Medical School, Boston, MA 02115, USA

^49^ Department of Evolution and Ecology and the Center for Population Biology, University of California, One Shields, Davis, CA 95616, USA

^50^ Department of Computer Science, Johns Hopkins University, Baltimore, MD 21218, USA

^51^ Department of Genetics, Epigenetics Institute, Perelman School of Medicine, University of Pennsylvania, Philadelphia, PA 19104, USA

^52^ Department of Pediatrics, Division of Genetics, School of Medicine, University of California, Irvine, CA 92697, USA

^53^ Sun Yat-sen University, Guangzhou, China

^54^ Edison Family Center for Genome Sciences & Systems Biology, Washington University School of Medicine, St. Louis, MO 63110, USA

^55^ Department of Biology and Center for Medical Genomics, Penn State University, University Park, PA 16802, USA

^56^ Division of Medical Genetics, Department of Medicine, University of Washington School of Medicine, Seattle, WA 98195, USA

^57^ The Jackson Laboratory for Genomic Medicine, Farmington, CT 06032, USA

^58^ Department of Biology, Penn State University, University Park, PA 16802, USA

^59^ Department of Biomedical Science, College of Health Sciences, Qatar University, Doha, Qatar

^60^ Department of Genetic Medicine, Weill Cornell Medicine-Qatar, Doha, Qatar

^61^ IRSD - Digestive Health Research Institute, University of Toulouse, INSERM, INRAE, ENVT, UPS, Toulouse, FR

^62^ MATCH biosystems, S.L., Elche, Spain

^63^ Universidad Miguel Hernández de Elche, Elche, Spain

^64^ Department of Computational Biology and Medical Sciences, The University of Tokyo, Kashiwa, Chiba 277-8561, Japan

^65^ Department of Computer Science, University of Pisa, Pisa, Italy

^66^ Law School, University of Wisconsin-Madison, Madison, WI 53706, USA

^67^ Institute of Genetics and Biomedical Research, UoS of Milan, National Research Council, Milan, Italy

^68^ Genome Biology Unit, European Molecular Biology Laboratory (EMBL), Heidelberg, DE

^69^ Institute for Molecular Medicine Finland, Helsinki Institute of Life Science, University of Helsinki, Helsinki, Finland

^70^ The Center for Bio- and Medical Technologies, Moscow, RUS

^71^ Centre for Biomedical Research and Technology, HSE University, Moscow, RUS

^72^ Department of Biology, Johns Hopkins University, Baltimore, MD 21218, USA

^73^ Coriell Institute for Medical Research, Camden, NJ 08103, USA

^74^ University of Amsterdam, Amsterdam, Netherlands

^75^ School of Clinical Medicine, University of Cambridge, Cambridge, CB2 0SP, UK

^76^ Center for Genomic Discovery, Mohammed Bin Rashid University, Dubai Health, UAE

^77^ Dubai Health Genomic Medicine Center, Dubai Health, UAE

^78^ GenomeArc Inc, Mississauga, ON, Canada

^79^ Department of Biology and Biotechnologies “Charles Darwin”, University of Rome “La Sapienza”, Rome 00185, IT

^80^ Center for Genomics, Loma Linda University School of Medicine, Loma Linda, CA 92350, USA

^81^ PacBio, Menlo Park, CA 94025, USA

^82^ The first affiliated hospital of Xi’an Jiaotong University, Xi’an Jiaotong University, Xi’an, Shaanxi, 710049, China

## Extended Data

**Fig. S1 | a,** PacBio HiFi read alignment depth from HG00099 aligning at the terminal ∼100kb of the chromosome in a donor-specific assembly (DSA, top), the CHM13 haploid assembly (middle), and HG002 diploid assembly (bottom) **b,** Proportion of primary alignments with MAPQ values above or equal to various thresholds in the terminal 100kb of the chromosome arms when aligning to DSA vs HG002 (diploid) and between the sequencing technologies used in this study (PacBio HiFi and ONT R10). **c,** NM tag value of reads (ONT R10 Fiber-seq reads, aligned to DSA) in the terminal 100kb of the chromosome arms, partitioned into reads overlapping the telomeres and those not overlapping the telomeres.

**Fig. S2 | a**, **f**, Histogram of the number of telomeric molecules per chromosome end in ONT R10 (**a**) and PacBio HiFi (**f**) samples. **b**, **g**, Number of haplotypes with an identified terminal telomeric repeat per chromosome arm for ONT R10 (**b**) and PacBio HiFi (**g**) datasets. **c**, **h**, Total number of telomeric long-read molecules per sample anchored at an identified subtelomere–telomere boundary in ONT R10 (**c**) and PacBio HiFi (**h**) datasets, per individual. **d**, **i**, Number of chromosome arm termini per sample with at least one molecule anchored at the subtelomere–telomere junction for ONT R10 (**d**) and PacBio HiFi (**i**) samples. **e**, **j**, Median number of anchored telomeric molecules per chromosome end, per individual, for ONT R10 (**e**) and PacBio HiFi (**j**) data.

**Fig. S3 | a**, Comparison of single-molecule TVR calls between ONT R10 and PacBio HiFi data. Single-molecule plots show TVRs ranging from 6 bp to 300 bp in size. **b**, TVRs across single molecules from all chr5q ends in the HPRC dataset (ONT R10). Each row represents a single read, with gray indicating TTAGGG repeats and color representing specific TVRs according to a shared color map. Data is filtered to include a minimum of 5 reads per individual haplotype end, with TVR sizes ranging from 1 to 300 bp. **c,** Distribution of TVR block sizes across all PacBio HiFi molecules comprising at least 1 kb of telomeric sequence. **d,** Boxplots displaying the maximum proportion of bases within individual TVRs covered by instances of a single *k*-mer. Coverage was evaluated across *k*-mer sizes ranging from 3 to 9 using ONT R10 and TVRs between 24 and 300 bp in length. **e**, Ranked bar plot representing the top 100 TVR 6-mers in PacBio (top) and ONT (bottom) data across all chromosome arms and individuals. **f,** TVRs as a proportion of the telomeric sequence per position for PacBio HiFi reads comprising at least 1 kb of telomeric sequence. Blue line uses all TVRs 1 - 300bp in size and the pink line is limited to TVRs 24 - 300bp in size to account for common, small PacBio sequencing errors in the telomere.

**Fig. S4 | a**, Four examples of TVR restructuring. Each row represents a single ONT R10 molecule. Colored bars indicate TVRs (colored by sequence content; Methods), and gray segments represent stretches of canonical TTAGGG repeats. All molecules are anchored at their respective subtelomere–telomere boundaries, with sequences shown in molecular coordinates.

**Fig. S5 | a**, Distribution of single-molecule telomere array lengths across HPRC samples in ONT R10 and PacBio HiFi reads overlapping the telomere-subtelomere junction. Minimum telomere array size 1kb.

**Fig. S6 | a**, Cumulative distribution of the TVR amount across molecules, partitioned by whether the corresponding chromosome arm is enriched for TGAGGG (arms 8q, 12q, 17p and Xp). Every faint line is an individual ONT R10 sample and the bold lines are the mean across all samples. **b,** Sorted bar plot of the 100 most common 6-mers in Xp, 12q and 8q across all individuals (ONT); TGAGGG in light blue all other 6-mers in gray. **c,** TVRs across single molecules from Xp, 8q and 12q chromosome arm ends across HPRC (ONT R10). Each row represents a single read, with gray indicating canonical TTAGGG repeats and color representing TVRs.

**Fig. S7 | a**, Subtelomeric TAR1 composition across all HPRC assemblies. Each horizontal bar represents the chromosome arm from a single haplotype, grouped by chromosome arm assignment. Green represents TAR1, gray is non-TAR1 subtelomeric sequence, and blue is telomeric sequence. Displaying 10kb of subtelomeric sequence. **b,** Bar plot of the proportion of contigs containing a TAR1 element within 10kb of the subtelomere-telomere boundary for each chromosome arm. **c,** Cumulative distribution of the TVR load across molecules per sample, stratified into arms with or without TAR1. Bold lines is the mean cumulative distribution across all PacBio samples.

**Fig. S8 | a,** m6A methylation events across single molecules at TAR1-containing chromosome arms in HG002 (top) and CHM13 (middle), compared to a TAR1-negative chromosome arm in CHM13 (bottom). **b,** Proportion of telomere-overlapping molecules containing a large MTase-accessible patch (>150 bp) across TAR1-positive (HG002, CHM13) and TAR1-negative (CHM13) chromosome arms. Points represent individual chromosome arms (*p* values determined by Mann–Whitney *U* tests). **c,** CHM13 molecules displayed as in **a**, with the addition of a CTCF motif and orientation track, and a zoomed-in view of the CTCF-bound region of a telomere proximal TAR1 element. **d,** Proportion of CHM13 PacBio Fiber-seq molecules with at least one footprinted CTCF binding site in the terminal 10 kb of the subtelomere versus regions 10 kb–1 Mb from the subtelomere–telomere boundary (Mann–Whitney *U* test). **e,** Distance between bound CTCF binding sites on Fiber-seq molecules containing multiple footprinted motifs, comparing the terminal 10 kb of the subtelomere to regions 10 kb–1 Mb from the subtelomere–telomere boundary ((Mann–Whitney *U* test). **f,** Proportion of CTCF motifs across all HPRC assemblies with the N-terminus oriented away from the telomere, at varying distances from the telomere (where *n* represents the total number of CTCF motifs). **g,** Number of CTCF motifs identified within each distance bin from the telomere across all chromosome arms in HPRC assemblies.

**Fig. S9 | a,** Comparison of single-molecule TVR calls across a trio. In each example, offspring is shown on top and parents below (HG002, son; HG003, father; HG004, mother). All reads are PacBio CCS reads from Genome in a Bottle (GIAB).

**Fig. S10 | a,** Single-molecule TVR plots for all p-arms with PacBio HiFi data in the HPRC dataset (excluding acrocentric chromosomes). Each row represents a single molecule, randomly selected for each haplotype, clustered based on their TVR similarity score in the proximal 200bp of the telomere.

**Fig. S11 | a,** Single-molecule TVR plots for all q-arms with PacBio HiFi data in the HPRC dataset (excluding acrocentric chromosomes). Each row represents a single molecule, randomly selected for each haplotype, clustered based on their TVR similarity score in the proximal 200bp of the telomere.

**Fig. S12 | a,** Two examples of TVR codes shared across multiple chromosome arms. Each cluster of rows represents individual molecules from a single haplotype and chromosome arm. Maximum of 10 molecules per arm; for arms exceeding this depth, a random subset of 10 molecules is displayed. Minimum TVR similarity score (proximal 200bp) of 0.7 used to identify clusters. **b,** SVbyEye visualization of an all-vs-all alignment of the 100-kb subtelomeric sequence proximal to the telomere in a subset of arms, including those highlighted in **a** (right, dashed box).

**Fig. S13 | a,** Single molecules from all 50 telomere arrays containing a detected TVR duplication event (boxed). For nested duplications, the first event is outlined with a dashed line and the second with a solid line. For chromosome arms with more than 10 molecules, 10 randomly selected molecules are displayed. For labels, “_1” denotes paternal and “_2” denotes maternal haplotypes. **b–d**, Zoomed-in view of the three telomere arrays exhibiting nested TVR duplication events.

**Fig. S14 | a,** Histogram of the number of telomeric molecules per chromosome end across HPRC ONT R10 Fiber-seq samples. **b**, Number of haplotypes with a molecule overlapping the subtelomere-telomere junction, per chromosome arm. **c,** Total number of telomeric Fiber-seq molecules. **d**, Number of chromosome arms per sample with at least one telomeric Fiber-seq molecule. **e**, Median number of telomeric Fiber-seq molecules per chromosome end, per individual.

**Fig. S15 | a**, Heatmaps of relative MTase-protected patch size distributions in relation to the subtelomere-telomere junction (in molecular coordinates) across 5 different HPRC Fiber-seq samples. MTase-protected patch sizes (y axis) and the positions they cover (x axis) are partitioned in 25-bp and 5-bp bins, respectively. **b**, Boxplots of the proportion of MTase protected patches larger than x bp at various values of x. The genome-wide distribution represents a 1% sample of all molecules across the genome, filtered to exclude those overlapping satellite elements. **c**, 2D histogram of telomere molecule length vs. terminal nucleosome position (MTase protected patch ≤210bp) per molecule. **d**, Cumulative fraction of the distances between MTase-protected patches of various sizes and of subtelomeric nucleosome size for reference. **e,** Boxplots displaying the mean fraction of bases within an MTase-accessible patch across the 0–2.5 kb and 2.5–5.0 kb regions of the telomere. Each data point represents an individual sample (Wilcoxon test *p*-value).

**Fig. S16 | a**, Density histogram of total methylation rate of all TVRs versus all TTAGGG sequences per molecule (log scale, Mann-Whitney U test) Faint lines represent individual samples, bold line is the mean distribution across samples. **b,** (Top) Relative methylation rates in the telomere for common TVR 6-mers. The methylation rate of each TVR was first normalized by it’s genome-wide methylation rate, and is expressed relative to the identically normalized canonical repeat. (Bottom) Number of TVR occurrences observed per individual. **c,** Per-individual skew of identified p- vs q- arms. **d,** Per-individual skew of the m6A-modified base per-read (A or T possible) in telomere-overlapping Fiber-seq reads.

